# High-resolution analysis of T cell subsets, clonality, and trajectories in psoriasis

**DOI:** 10.64898/2026.10.23.761402

**Authors:** Xin Luo, Jared H. Liu, Dev Bhatt, Joy Q. Jin, David Wu, Samuel Alber, Marwa Hakimi, Jinyeong Lim, Faye Orcales, Samuel Yeroushalmi, Zhi-Ming Huang, Tina Bhutani, Paolo Manzanillo, Wilson Liao

## Abstract

Psoriasis is sustained by type 17 immunity, yet how transcriptionally defined T-cell states are clonally organized across lesions, clinically uninvolved skin, and the circulation remains incompletely understood. We performed paired single-cell RNA and T-cell receptor (TCR) sequencing of T cells isolated from two spatially separated lesions, non-lesional skin, and peripheral blood from 23 participants with psoriasis, together with healthy skin and blood comparators. We resolved 19 cutaneous T-cell populations and found that lesions were enriched for IL-17-associated CD4+ and CD8+ states and activated regulatory T-cell states. CD8+ Tc17 cells showed prominent clonal expansion, and expanded clonotypes contributed more prominently to skin than to blood. Clonotype sharing was greatest between paired lesions, while clonotypes detected in cutaneous IL-17-associated cells mapped to multiple circulating phenotypes, indicating that blood and skin capture complementary features of the disease-associated repertoire. At the sample level, T-cell composition separated 57 lesional samples into two recurrent compositional states that were largely concordant across paired body sites. HLA-C*06:02 status was associated with differences in pretreatment lesional CD8+ T-cell composition, clonotype overlap, and clonal-size distributions. Exploratory longitudinal analyses in eight treated participants showed heterogeneous remodeling of T-cell composition and clonotype detection after therapy. Together, these findings define tissue-restricted and tissue-shared features of the psoriatic T-cell repertoire and identify clonal architecture across skin sites and blood as an organizing feature of lesion heterogeneity.

## Introduction

Psoriasis is a chronic immune-mediated skin disease in which genetic susceptibility, cutaneous antigen recognition, and cytokine signaling converge to sustain tissue inflammation. T cells are central to this process: psoriasis risk loci are enriched in pathways involving antigen presentation and T-cell activation, HLA-C*06:02 is the strongest common genetic association, and the clinical efficacy of therapies targeting the IL-23–IL-17 axis demonstrates the importance of type 17 immunity ^1,2^. Candidate autoantigens, including LL37 and melanocyte-derived peptides, further support a role for antigen-experienced T cells in at least a subset of lesions, although the antigenic determinants and relevant T-cell populations remain incompletely defined ^3,4^.

Studies of human skin have revealed marked functional and transcriptional heterogeneity among cutaneous T cells. Work from Liao and colleagues identified distinct IL-17A-expressing CD8+ T-cell states in psoriatic skin and showed that Tc17 cells coexist with other inflammatory, resident, and recirculating CD8+ populations ^5^. Complementary single-cell analyses at UCSF described a transcriptional gradient linking cutaneous memory T-cell states while restricting Th17 and Tc17 identities ^6^. Other studies have shown that IL-17-producing alpha-beta T-cell clones can persist in clinically resolved lesions, consistent with a durable local immune memory ^7^. Cutaneous regulatory T cells (Tregs) are similarly heterogeneous. Joint work from the Liao and Rosenblum laboratories identified CD27- and OX40-dependent constraints on Th17-like differentiation in Tregs, providing a mechanistic framework for interpreting activated Treg states without equating marker expression with suppressive capacity ^8^. Longitudinal profiling from the same UCSF collaborative environment further showed that IL-17A blockade remodels cutaneous Treg as well as effector T-cell transcriptomes ^9^.

Despite these advances, the relationships among T-cell state, clonal expansion, and tissue distribution in psoriasis remain incompletely resolved. Prior studies have often emphasized a single tissue compartment or T-cell lineage, and transcriptomic and TCR analyses have not always been integrated across matched samples. It is therefore unclear how transcriptionally defined T-cell states are clonally organized across paired lesions, clinically uninvolved skin, and blood; whether lesions from the same individual share a reproducible T-cell composition; and whether HLA-C*06:02 status is associated with this cellular and clonal architecture. These questions are particularly important because comparisons of circulating and tissue-resident immune populations can reveal disease-associated clonotypes that are not apparent from either compartment alone ^10^.

Here, we performed paired single-cell RNA and TCR sequencing of T cells isolated from two psoriasis lesions, non-lesional skin, and peripheral blood from the same participants. We defined the major cutaneous CD4+ and CD8+ states, quantified their clonal expansion, and mapped clonotype sharing across tissues and transcriptional phenotypes. Lesional skin was enriched for IL-17-associated CD4+ and CD8+ states and activated Treg states, while expanded clonotypes contributed more prominently to skin than to blood. Clonotype sharing was greatest between paired lesions and connected cutaneous IL-17-associated cells to multiple circulating phenotypes. At the sample level, T-cell composition identified two recurrent lesional compositional states that were largely concordant across paired body sites. We also observed associations between HLA-C*06:02 status and CD8+ T-cell composition and clonal organization, and we describe exploratory longitudinal observations after treatment. Together, these data provide an integrated view of how T-cell phenotype, clonality, and tissue distribution are organized in psoriasis.

## Results

### Single-cell profiling defines the T-cell landscape of psoriatic skin

We isolated CD45+CD3+ T cells from lesional, contralesional, and non-lesional skin biopsies and peripheral blood mononuclear cells (PBMCs) from patients with psoriasis and performed paired single-cell RNA sequencing and T-cell receptor (TCR) sequencing using the 10x Genomics 5′ platform. Two psoriasis lesions on opposite sides of the body (termed “lesional” and “contralesional”) and one non-lesional skin site were sampled per patient. We retained cells that passed mitochondrial, gene-expression, and RNA-content quality filters and contained one productive paired TCR alpha and beta chain (Fig. S1). Ambiguous cell clusters were assigned to CD4+ or CD8+ compartments using CITE-seq protein tags or shared TCR clonotype identity with cells in unambiguous clusters. Sample-level cohort, tissue, and longitudinal metadata are provided in Table 1.

A total of 19 transcriptionally distinct skin-derived T-cell clusters were identified based on canonical phenotypic markers (Fig. 1A–B). Cluster-composition analysis identified multiple populations with significantly different relative abundance in lesional and non-lesional skin (Fig. 1C, Fig. S2C). A cluster containing IL-17-producing CD4+ and CD8+ T cells was strongly enriched in lesional skin. CD4+ regulatory T cells (Tregs) were also more frequent in lesional skin, whereas CD8+ effector-memory and CD4+ CD44+ effector-memory populations were relatively enriched in non-lesional skin (Fig. 1C). In paired samples, the increase in IL-17-producing T cells exceeded the increase in Tregs, resulting in a higher T17/Treg ratio in lesional skin (Fig. 1D).

**Figure 1.**
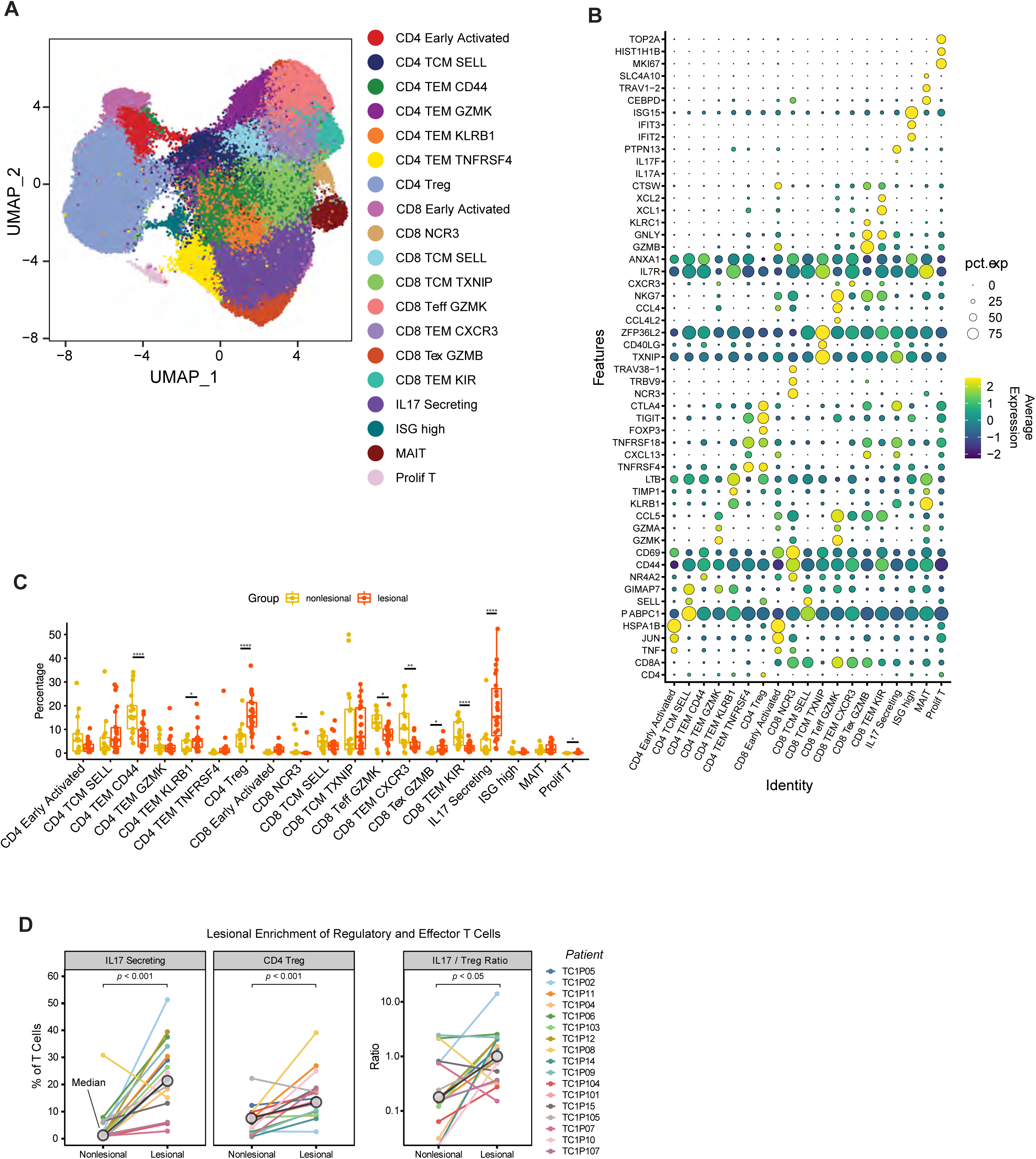
Single-cell profiling defines the T-cell landscape of psoriatic skin. (A) Uniform manifold approximation and projection (UMAP) of skin-derived T cells in the global analysis, colored by 19 annotated T-cell states. (B) Dot plot of marker-gene expression across the global states. Dot size indicates the percentage of cells expressing each gene, and color indicates scaled average expression. (C) Sample-level frequencies of the annotated states in pretreatment lesional (n = 23 samples) and non-lesional (n = 17 samples) psoriasis skin. Each point represents one sample; boxes summarize the median and interquartile range. Differential abundance was evaluated with DCATS with false-discovery-rate (FDR) correction; one or more asterisks indicate FDR < 0.05. (D) Paired frequencies of IL-17-secreting T cells and CD4+ regulatory T cells (Tregs), and the IL-17-secreting T-cell/Treg ratio, in matched lesional and non-lesional samples (n = 17 participants). Colored lines connect samples from the same participant, gray circles indicate group medians, and the ratio is shown on a logarithmic scale. All three displayed paired comparisons have P < 0.001.

### Lesional skin is enriched for CD8+ Tc17 cells

IL-17-producing CD8+ T cells have previously been identified in psoriasis lesions by flow-cytometric and single-cell studies ^5,11^. To define CD8+ T-cell heterogeneity in our cohort, we selected CD8+ T cells with paired TCR sequences and reclustered them into 16 transcriptional states (Fig. 2A–B). These included multiple effector-memory populations with distinct transcriptional profiles. Tc17 cells were strongly enriched in lesional skin, whereas CD44+ effector-memory cells were relatively enriched in non-lesional and healthy skin (Fig. 2C). Cell-density distributions similarly shifted from CD44+ effector-memory states toward Tc17 cells in lesional samples (Fig. 2D). RNA-velocity vectors were consistent with trajectories connecting early activated and central- and effector-memory states with Tc17 cells; these computational relationships do not by themselves establish lineage direction or a terminal differentiation state (Fig. 2E).

**Figure 2.**
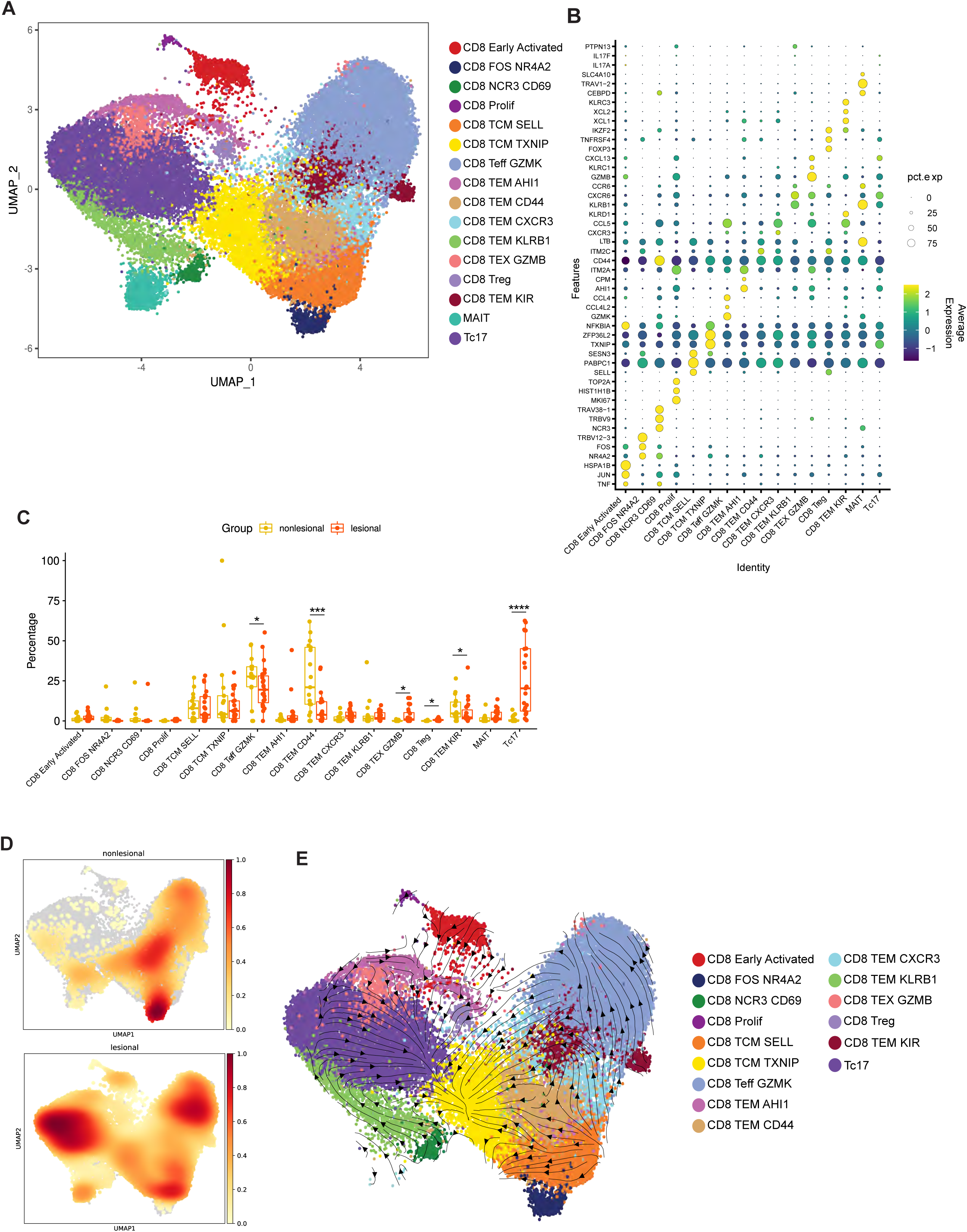
Phenotypic heterogeneity of CD8+ T cells in psoriasis skin. (A) UMAP of skin-derived CD8+ T cells, colored by 16 annotated states. (B) Dot plot of marker-gene expression across CD8+ states; dot size indicates the percentage of cells expressing each gene, and color indicates scaled average expression. (C) Sample-level frequencies of CD8+ states in pretreatment lesional (n = 23 samples) and non-lesional (n = 17 samples) psoriasis skin. Each point represents one sample; boxes summarize the median and interquartile range. Differential abundance was evaluated with DCATS with FDR correction; one or more asterisks indicate FDR < 0.05. (D) Gaussian-kernel density of CD8+ cells across the UMAP in non-lesional (top) and lesional (bottom) samples. Density values were scaled from 0 to 1 within each group. (E) RNA-velocity streamlines projected onto the CD8+ UMAP. Mucosal-associated invariant T (MAIT) cells were excluded from the velocity visualization; streamlines summarize transcriptional dynamics and do not establish lineage direction.

### CD4+ effector and regulatory T-cell states are remodeled in lesional skin

CD4+ IL-17-producing and regulatory T-cell populations have both been implicated in psoriatic inflammation, and previous studies have described altered Treg phenotype and function in psoriasis ^8,12,13^. We therefore analyzed conventional CD4+ T cells and Tregs separately. Reclustering of conventional CD4+ T cells resolved distinct central-memory, effector-memory, early activated, and Th17-associated populations, with identities assigned by canonical marker expression (Fig. 3A–B).

**Figure 3.**
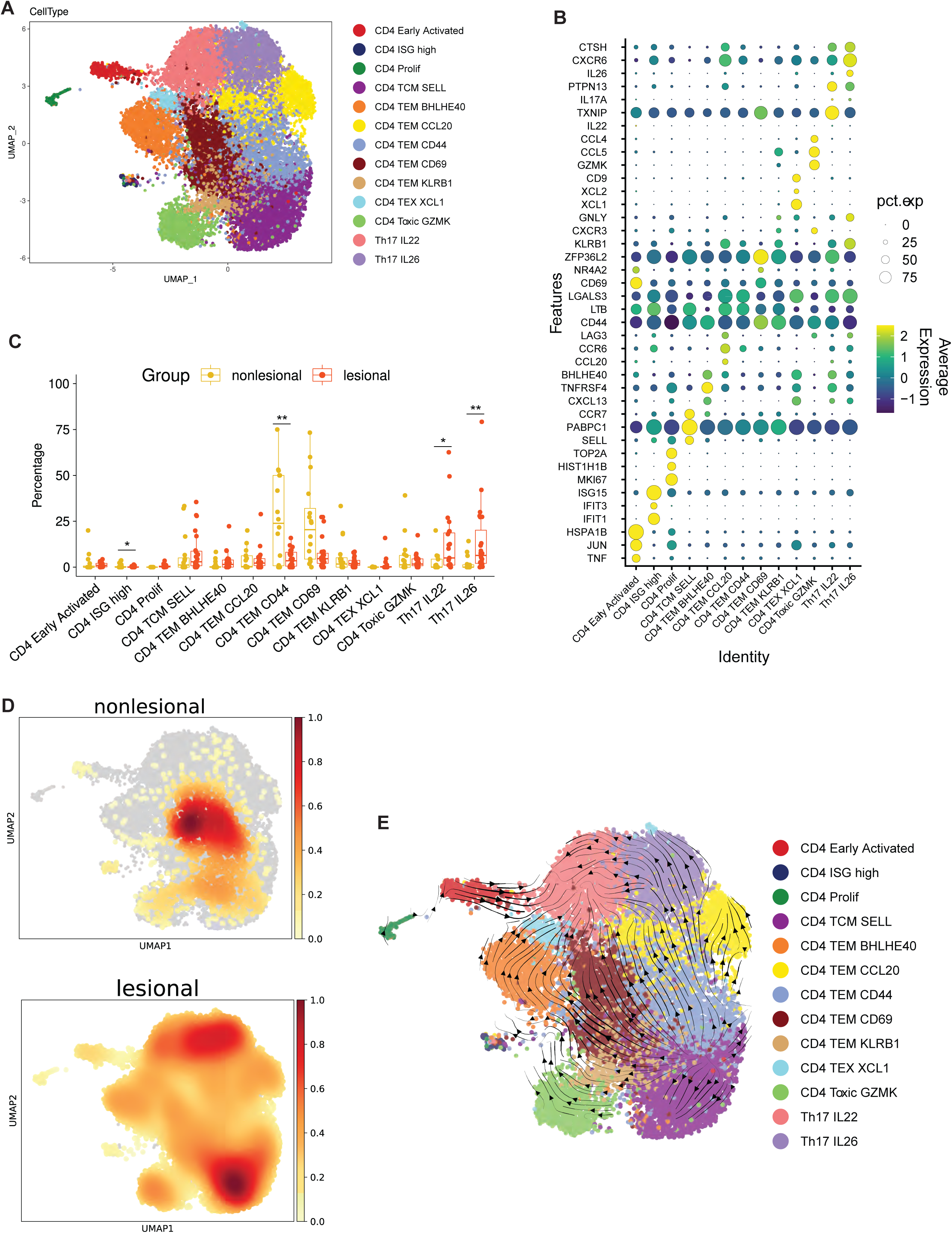
Phenotypic heterogeneity of conventional CD4+ T cells in psoriasis skin. (A) UMAP of skin-derived conventional CD4+ T cells after exclusion of Tregs, colored by 13 annotated states. (B) Dot plot of marker-gene expression across conventional CD4+ states; dot size indicates the percentage of cells expressing each gene, and color indicates scaled average expression. (C) Sample-level frequencies of conventional CD4+ states in pretreatment lesional (n = 23 samples) and non-lesional (n = 16 samples) psoriasis skin. Each point represents one sample; boxes summarize the median and interquartile range. Differential abundance was evaluated with DCATS with FDR correction; one or more asterisks indicate FDR < 0.05. (D) Gaussian-kernel density of conventional CD4+ T cells across the UMAP in non-lesional (top) and lesional (bottom) samples, scaled from 0 to 1 within each group. (E) RNA-velocity streamlines projected onto the conventional CD4+ UMAP. Streamlines summarize transcriptional dynamics and do not establish lineage direction.

The relative abundance of conventional CD4+ T-cell states differed between lesional and non-lesional skin (Fig. 3C). Th17-associated populations were increased in lesional samples, accompanied by a relative reduction in CD44+ effector-memory cells. Cell-density distributions showed the same shift toward IL-17-secreting and SELL+ central-memory states in lesional skin (Fig. 3D). RNA-velocity vectors connected effector-memory, early activated, and Th17-associated states, consistent with more than one potential transcriptional route into the lesional Th17 compartment; this analysis does not establish direct differentiation between these populations (Fig. 3E).

Because Tregs were more abundant in lesional than non-lesional skin, we further characterized their transcriptional heterogeneity (Fig. 4A–D). Lesional Tregs were shifted away from memory-associated states toward states with higher expression of TIGIT, CTLA4, ICOS, and related activation-associated markers. These marker patterns are consistent with activated regulatory phenotypes but do not establish greater suppressive function. RNA-velocity analysis suggested transcriptional relationships among the Treg states without resolving a definitive lineage sequence (Fig. 4E).

**Figure 4.**
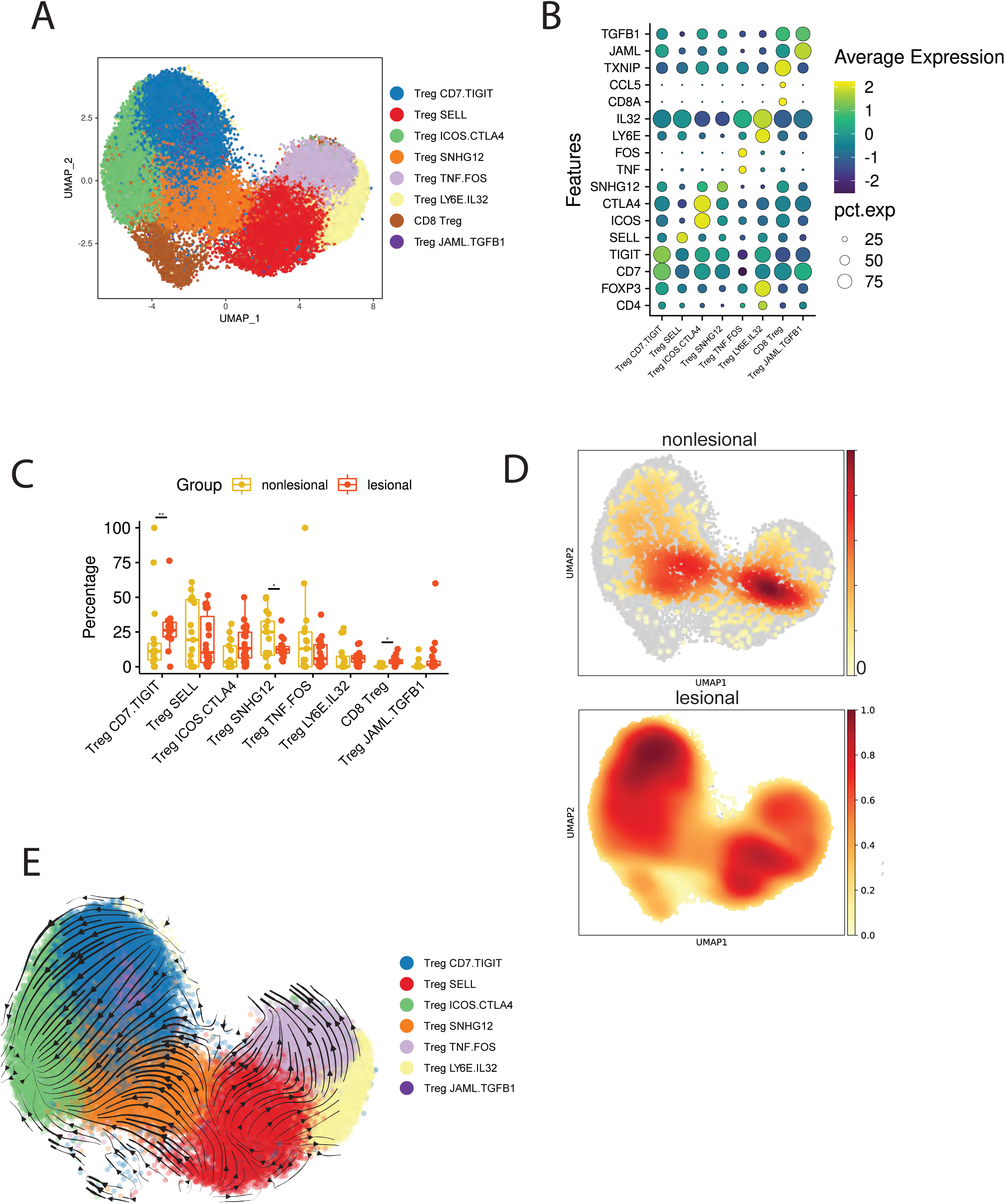
Transcriptional heterogeneity of regulatory T cells in psoriasis skin. (A) UMAP of regulatory T-cell subclusters, colored by annotated state. (B) Dot plot of marker-gene expression across regulatory T-cell states; dot size indicates the percentage of cells expressing each gene, and color indicates scaled average expression. (C) Sample-level frequencies of regulatory T-cell states in pretreatment lesional (n = 23 samples) and non-lesional (n = 17 samples) psoriasis skin. Each point represents one sample; boxes summarize the median and interquartile range. Differential abundance was evaluated with DCATS with FDR correction; one or more asterisks indicate FDR < 0.05. (D) Gaussian-kernel density of regulatory T cells across the UMAP in non-lesional (top) and lesional (bottom) samples, scaled from 0 to 1 within each group. (E) RNA-velocity streamlines projected onto the CD4+ Treg states; the CD8 Treg state is not displayed in this velocity panel. Streamlines summarize transcriptional dynamics and do not establish lineage direction.

### Skin and blood share major T-cell phenotypes but differ in composition

To examine relationships between skin and blood T cells, we integrated transcriptomic profiles from both compartments. All but one of the 22 phenotypic populations identified in the integrated analysis contained cells from both skin and blood (Fig. 5A–C); CD8+ NCR3 T cells were detected only in skin from a single patient. Several populations were disproportionately represented in blood, including CD4+ naïve and CD4+ TCM SELL cells, whereas IL-17-secreting and early activated T cells were disproportionately represented in skin (Fig. 5B–C).

**Figure 5.**
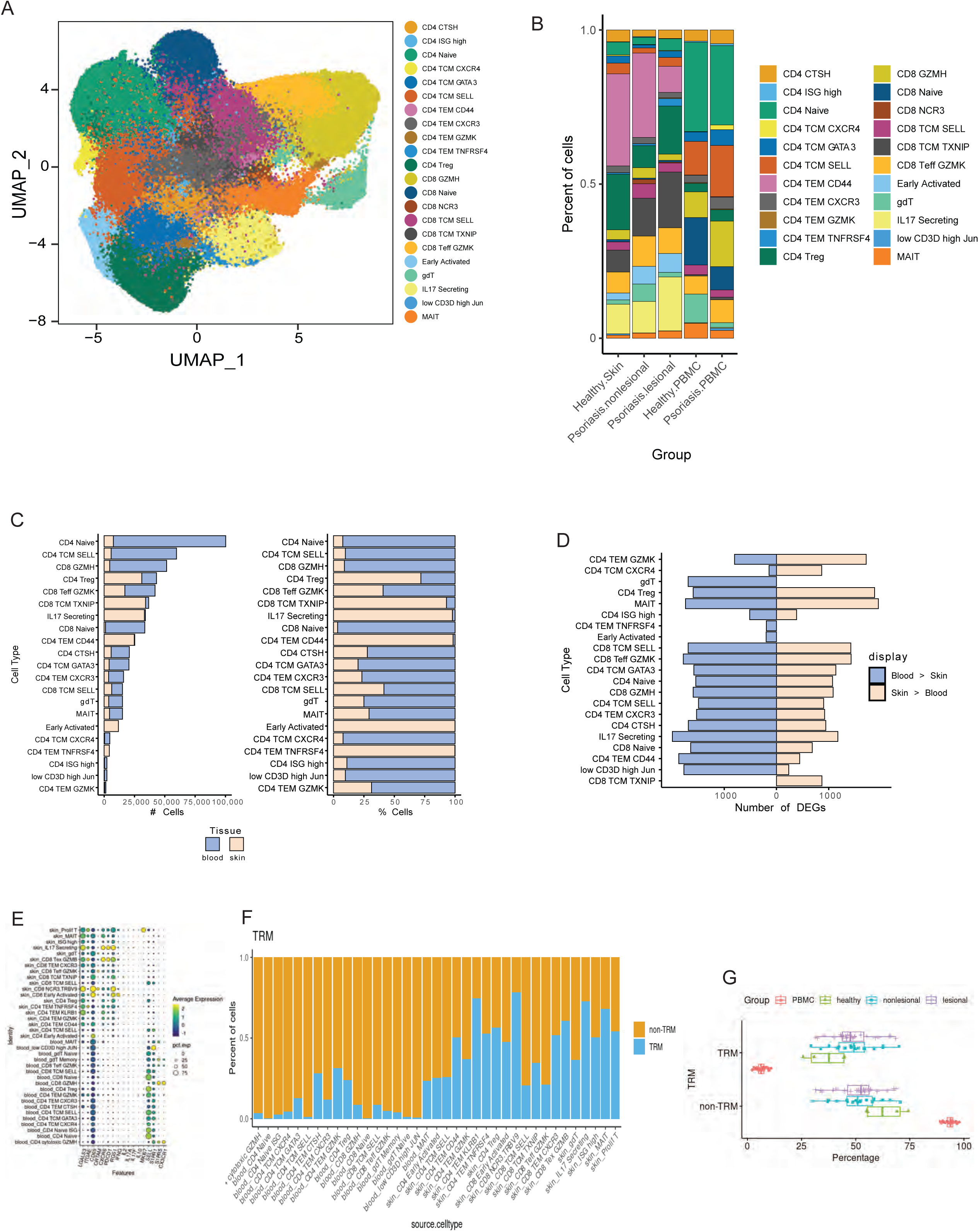
Integrated analysis of skin and blood T cells. (A) UMAP of the integrated skin and peripheral blood mononuclear cell (PBMC) dataset from participants with psoriasis and healthy donors, colored by 22 annotated T-cell populations. (B) Relative abundance of the annotated populations in healthy skin, non-lesional psoriasis skin, lesional psoriasis skin, healthy PBMCs, and psoriasis PBMCs. (C) Total number of cells from skin and blood in each population (left) and the percentage contribution of each tissue compartment within each population (right). (D) Number of genes called differentially expressed between blood and skin within each annotated population, separated by direction of effect (blue, higher in blood; beige, higher in skin). These bars summarize the number and direction of differentially expressed genes, not their effect sizes. (E) Dot plot of selected tissue-residency- and recirculation-associated genes across source-specific populations. Dot size indicates the percentage of cells expressing each gene, and color indicates scaled average expression. (F) Proportions of cells classified as TRM-like or non-TRM-like within each source-specific population using the study’s transcriptional signature criteria. (G) Sample-level percentages of TRM-like and non-TRM-like cells in PBMC, healthy-skin, non-lesional psoriasis-skin, and lesional psoriasis-skin groups. Points represent samples; boxes summarize the median and interquartile range.

At the level of aggregate composition, most major T-cell populations were present at broadly similar frequencies in healthy and psoriasis PBMC samples. Psoriasis PBMC samples showed descriptive shifts toward GZMH+ effector cells and away from naïve T cells, but these modest differences were not a dominant feature of the integrated analysis (Fig. 5B).

### Skin T cells exhibit tissue-residency-associated transcriptional programs

Human tissue-resident memory T cells (TRM) are characterized by increased expression of tissue-retention and adhesion programs, including ITGAE and CXCR6, and reduced expression of recirculation-associated genes such as S1PR1 and SELL ^14^. We evaluated these programs across skin and blood populations (Fig. 5E–G). Most skin T-cell clusters expressed TRM-associated genes, whereas blood-derived populations expressed higher levels of recirculation-associated markers (Fig. 5E). The fraction of cells meeting the study’s TRM-signature criteria was higher in both lesional and non-lesional psoriasis skin than in healthy skin, with no clear difference between the two psoriasis skin sites (Fig. 5F–G). This pattern is consistent with prior observations of persistent IL-17-producing resident T-cell populations in psoriasis skin ^7,15^.

### Lesional and non-lesional skin show distinct patterns of clonal expansion

We next analyzed paired TCR alpha- and beta-chain sequences to quantify clonal expansion in blood and skin. Clonotypes were defined by exact matches of paired alpha- and beta-chain sequences and grouped as single, small, medium, large, or hyperexpanded according to the number of cells bearing each clonotype (Fig. 6A). Singletons accounted for 74.0% of blood T cells and 53.8% of skin T cells, indicating a larger contribution of expanded clonotypes in skin. Cluster-level distributions also differed between lesional and non-lesional skin (Fig. 6B). Large and hyperexpanded clonotypes were prominent within the lesional IL-17-secreting population, whereas several CD8+ central-memory and effector populations contained expanded clonotypes in non-lesional skin. These results identify compartment- and state-associated differences in clonal expansion without assigning antigen specificity to the expanded clonotypes.

**Figure 6.**
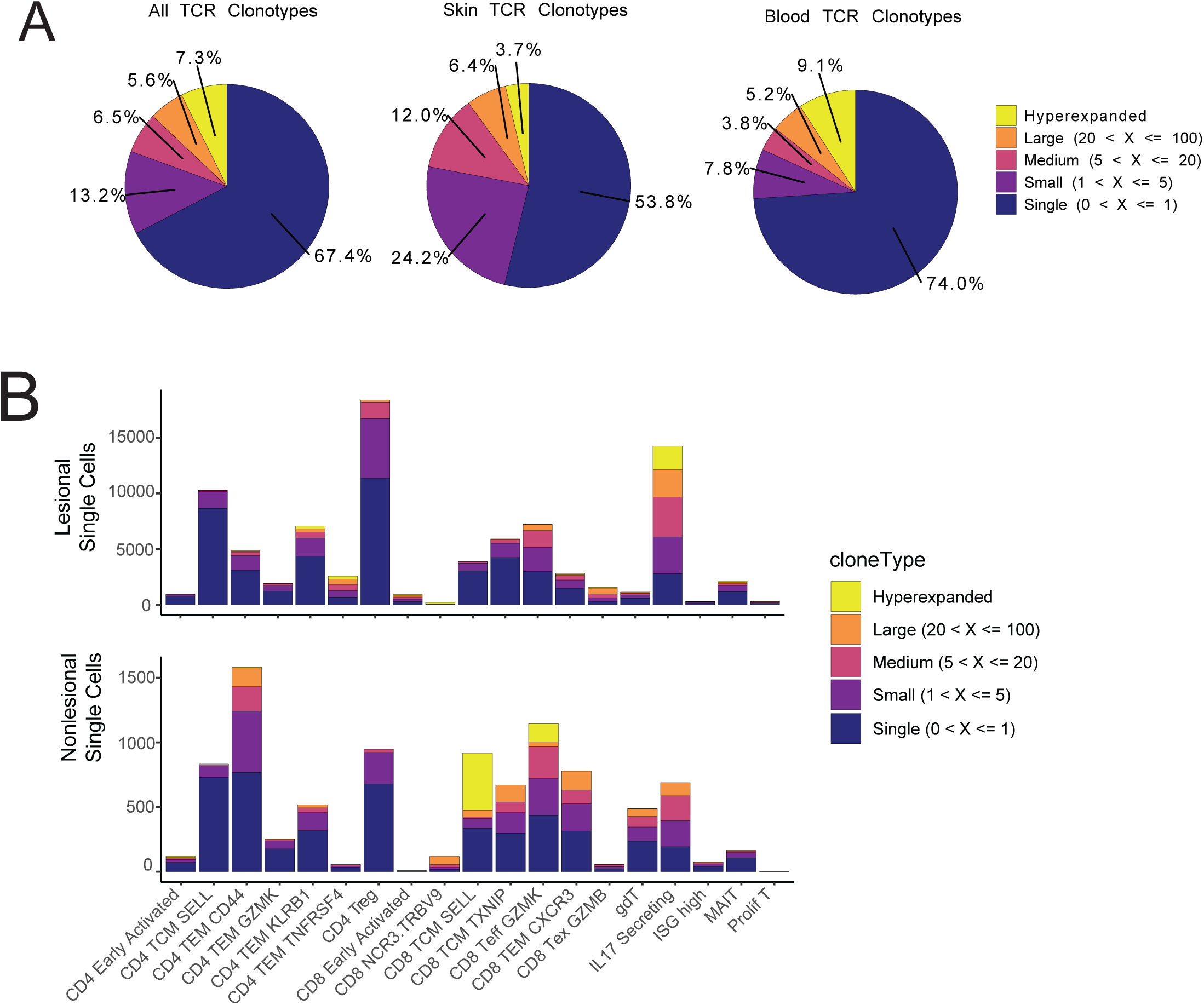
Clonal expansion differs between skin and blood. (A) Proportions of cells with one productive paired T-cell receptor (TCR) α/β clonotype assigned to each clonal-size category in the combined dataset (left), skin (middle), and blood (right). Categories were defined by the number of cells bearing the same clonotype: single, 1 cell; small, 2–5 cells; medium, 6–20 cells; large, 21–100 cells; and hyperexpanded, >100 cells. (B) Absolute numbers of cells with paired TCRs in each global skin T-cell state in lesional (top) and non-lesional (bottom) samples, stacked by clonal-size category.

### TCR clonotypes are shared across skin sites and, less frequently, with blood

We next quantified clonotype overlap among lesional, contralesional, and non-lesional skin and PBMCs (Fig. 7A–B). The greatest tissue-level overlap occurred between lesional and contralesional sites (Morisita index, 0.625), whereas overlap involving non-lesional skin or PBMCs was substantially lower (Fig. 7A). At the cell-state level, Tc17 clonotypes overlapped with several early activated, proliferating, and memory CD8+ states (Fig. 7C). CD4+ states showed a different and generally more limited pattern of pairwise clonotype overlap (Fig. 7D). These analyses demonstrate sharing among phenotypic states but do not establish precursor–product relationships.

**Figure 7.**
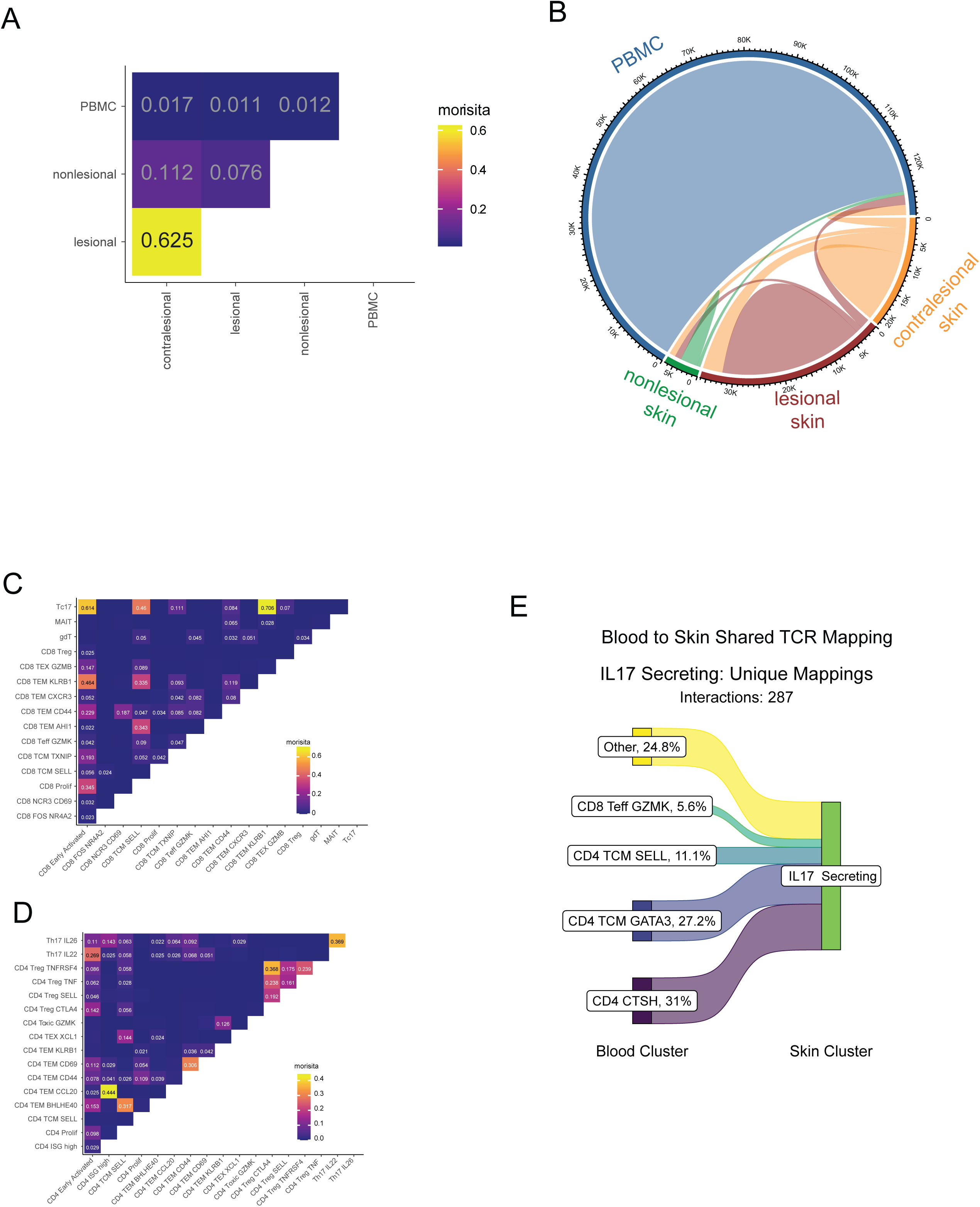
Clonotype sharing across tissues and T-cell states. (A) Pairwise Morisita overlap indices for paired TCR clonotypes among contralesional, lesional, and non-lesional skin and PBMCs. (B) Chord diagram of clonotypes shared between tissue compartments; ribbon width represents the extent of sharing. (C, D) Pairwise Morisita overlap among skin CD8+ states (C) and skin CD4+ states (D). Values >0.02 are displayed for clarity. (E) Blood-to-skin mapping of 287 unique cluster mappings involving paired TCR clonotypes detected in the skin IL-17-secreting population. Ribbons and labels show the percentage mapping to the four most frequent annotated blood populations; the remaining blood populations are combined as Other.

Although blood–skin overlap was limited relative to overlap between skin sites, shared clonotypes were detected across multiple blood and skin populations (Fig. 7A–E). Among 287 blood-to-skin mappings involving IL-17-secreting skin cells, the largest annotated blood populations were CD4+ CTSH cells (31.0%), CD4+ TCM GATA3 cells (27.2%), CD4+ TCM SELL cells (11.1%), and CD8+ Teff GZMK cells (5.6%); the remaining 24.8% were distributed across other populations (Fig. 7E). Thus, the cutaneous IL-17-secreting state was linked by shared TCR sequences to multiple circulating phenotypes rather than to a single blood population.

### Exploratory longitudinal analysis identifies heterogeneous changes after treatment

Eight patients contributed skin samples before and after heterogeneous biologic or phototherapy treatments. This longitudinal analysis was therefore considered exploratory. Most major transcriptional states remained detectable after treatment, although their relative composition varied among patients (Fig. 8A). Median frequencies of IL-17-secreting cells and Tregs decreased after treatment, but individual responses and the IL-17/Treg ratio were variable (Fig. 8B). Clonotype tracking showed both persistent and time-point-specific clonotypes, again with substantial interindividual variation (Fig. 8C). These observations describe within-patient changes but do not support treatment-specific mechanistic conclusions.

**Figure 8.**
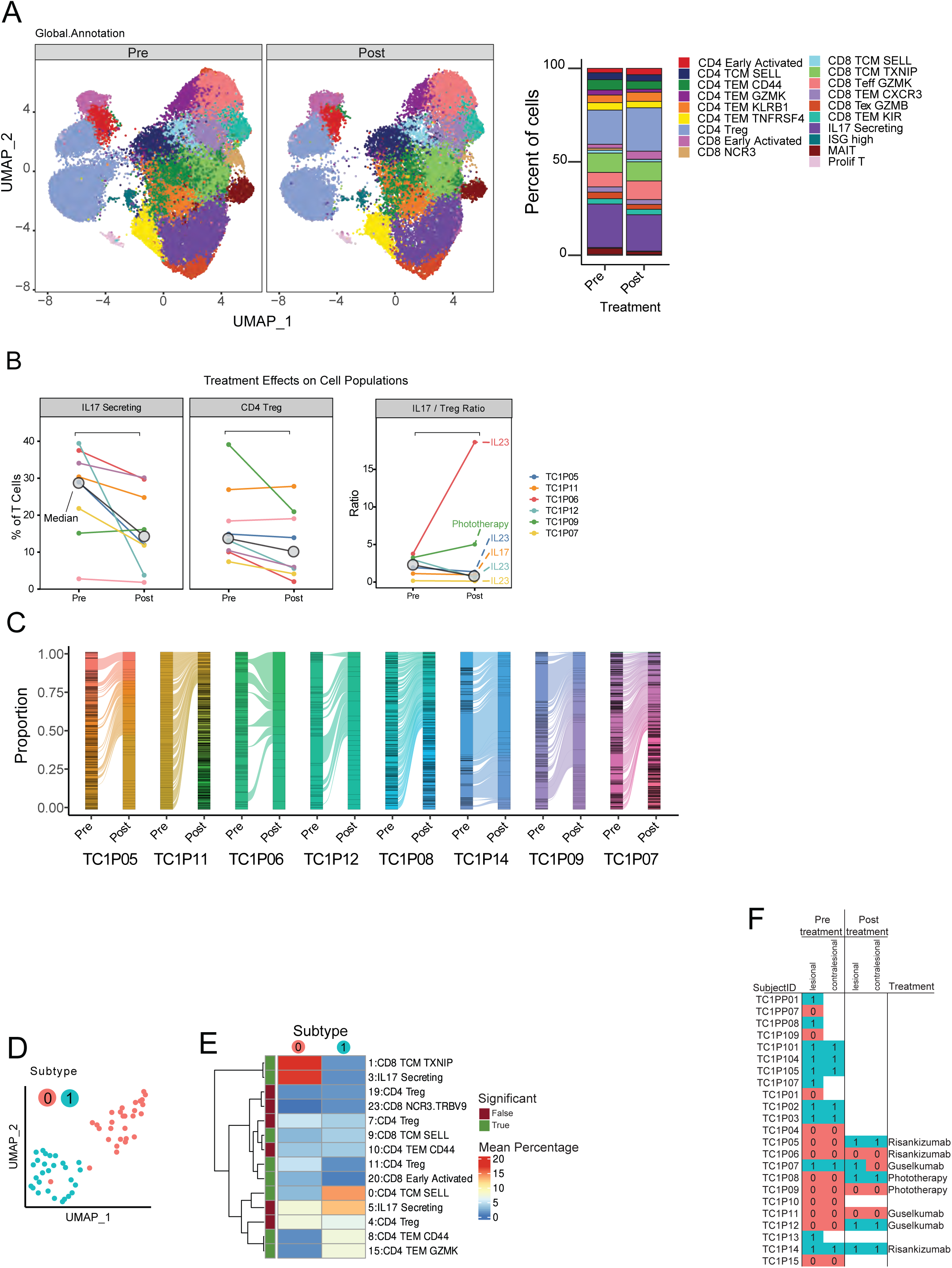
Exploratory longitudinal treatment analysis and lesional T-cell compositional states. (A) Global skin T-cell UMAPs split by pretreatment and post-treatment sampling (left) and aggregate cell-state composition at the two time points (right) for the longitudinal cohort (n = 8 participants: risankizumab, n = 3; guselkumab, n = 3; phototherapy, n = 2). (B) Participant-level pretreatment and post-treatment frequencies of IL-17-secreting T cells and CD4+ Tregs, and the corresponding IL-17/Treg ratio, for participants with paired composition estimates displayed (n = 6). Lines connect time points from the same participant, and gray circles indicate group medians. (C) Participant-level alluvial plots of paired TCR clonotypes before and after treatment (n = 8 participants). Ribbons connect clonotypes detected at both time points; unlinked segments represent time-point-restricted clonotypes, and each time-point bar is normalized to a total proportion of 1. (D) UMAP of percentile-ranked 24-cluster composition profiles from 57 lesional samples, colored by compositional state 0 (n = 29 samples) or state 1 (n = 28 samples). (E) Mean abundance of the 14 clusters contributing most strongly to separation of the two compositional states. The side annotation identifies clusters meeting the analysis significance criterion for a between-state difference. (F) Compositional-state assignments by participant, skin site, treatment time point, and treatment; blank cells indicate that no state is displayed for that participant/site/time point. Longitudinal comparisons are descriptive because the cohort was small and treatments were heterogeneous.

### T-cell composition identifies two recurrent lesional compositional states

Prior transcriptomic and spatial studies have described molecular heterogeneity among psoriasis lesions ^16-18^. We therefore asked whether lesional samples in our single-cell dataset could be grouped by T-cell composition. Using an archetype-style approach adapted from analysis of tumor immune ecosystems ^19^, we percentile-ranked the abundance of 24 de novo T-cell clusters across 57 lesional samples from 23 patients and identified 14 clusters that contributed most strongly to sample separation (Fig. 8D–E).

The 14-cluster composition profiles separated the lesional samples into two groups of similar size (state 0, n = 29; state 1, n = 28) (Fig. 8D–F). State 0 contained higher relative abundance of selected central-memory and early activated CD8+ populations, IL-17-secreting T cells, and CD4+ Tregs. State 1 was enriched for selected CD4+ effector-memory populations and different CD4+ and CD8+ central-memory populations (Fig. 8E). Because these groups were derived from T-cell composition and were not linked here to a validated clinical endpoint, we refer to them as compositional states rather than clinical disease subtypes.

### Lesional compositional states are largely concordant across paired body sites and may change over time

Lesional and contralesional biopsies from opposite body sites had concordant compositional-state assignments in all but one paired comparison (Fig. 8F). Among the eight patients sampled before and after treatment, three changed state assignment, whereas the others retained their initial assignment or had an ambiguous post-treatment classification (Fig. 8F). These observations suggest within-patient concordance across sampled lesions but provide only preliminary evidence about temporal stability because of the small, treatment-heterogeneous longitudinal cohort.

### HLA-C*06:02 status is associated with CD8+ T-cell composition and clonal architecture

HLA-C*06:02 is a major psoriasis susceptibility allele that shapes the class I peptide repertoire available for CD8+ T-cell recognition ^2,20,21^. We therefore compared CD8+ T-cell composition between HLA-C*06:02 carriers and non-carriers. CD8+ TCM SELL and CD8+ TEM CD44 populations were more frequent in carriers, whereas CD8+ early activated cells were more frequent in non-carriers. MAIT cells were also more frequent in carriers (Fig. 9A).

**Figure 9.**
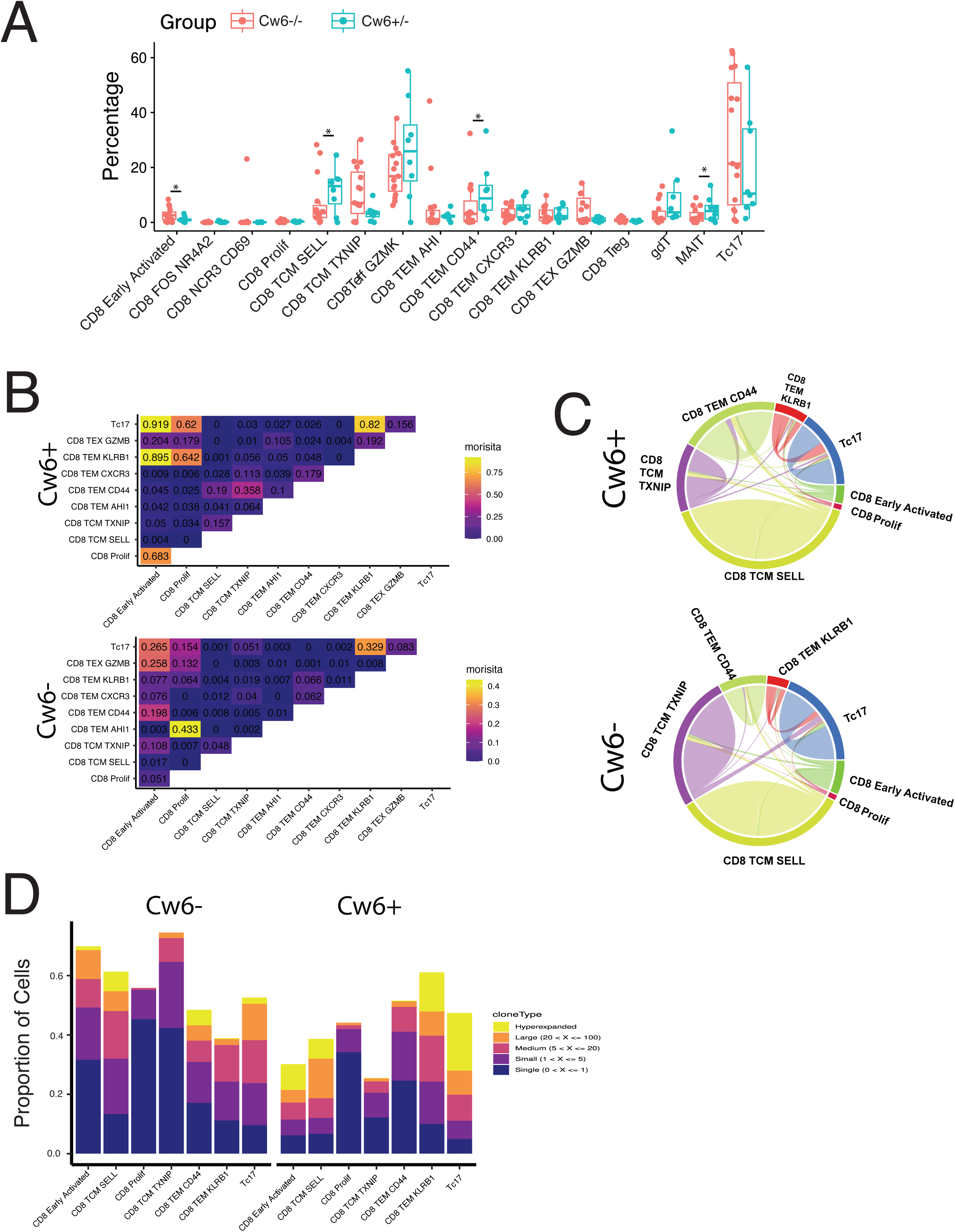
HLA-C*06:02 status is associated with pretreatment lesional CD8+ T-cell composition and clonal architecture. Participants carrying at least one HLA-C*06:02 allele are denoted HLA-C*06:02+ (n = 8), and non-carriers are denoted HLA-C*06:02− (n = 15). (A) Sample-level frequencies of CD8+ T-cell states by HLA-C*06:02 status. Each point represents one participant; boxes summarize the median and interquartile range. Differential abundance was evaluated with DCATS with FDR correction; *FDR < 0.05. (B) Pairwise Morisita overlap among selected CD8+ states in carriers (top) and non-carriers (bottom). (C) Chord diagrams summarizing clonotype sharing among selected CD8+ states in carriers (top) and non-carriers (bottom); ribbon width represents the extent of sharing. (D) Proportions of cells in selected CD8+ states assigned to each clonal-size category, stratified by HLA-C*06:02 status. Clonal-size categories are defined as in Figure 6.

Pairwise clonotype-overlap patterns also differed between genotype groups (Fig. 9B–C). In carriers, Tc17 cells showed greater overlap with CD8+ early activated, proliferating, and TEM KLRB1 cells than was observed in non-carriers. Clonal-size distributions also differed descriptively across selected CD8+ states, with conspicuous contributions from large or hyperexpanded clonotypes in carrier Tc17 and TEM KLRB1 populations (Fig. 9D). Because these comparisons are observational and formal between-genotype testing is not displayed for all clonotype metrics, they should be interpreted as genotype-associated patterns and do not establish antigen specificity or a causal effect of HLA-C*06:02 on clonal architecture. Additional descriptive comparisons of TCR variable-gene usage and conventional CD4+ and regulatory T-cell composition by HLA-C*06:02 status are provided in Fig. S3.

## Discussion

In this study, paired single-cell transcriptomic and TCR profiling across multiple skin sites and blood resolved several complementary dimensions of the T-cell response in psoriasis. Lesional skin was enriched for IL-17-associated CD4+ and CD8+ states as well as activated Treg states; expanded clonotypes contributed more strongly to skin than to blood; and clonotype overlap was greatest between paired lesions from the same participant. At the lesion level, T-cell composition identified two recurrent compositional states that were usually concordant across paired body sites. HLA-C*06:02 status was associated with differences in CD8+ T-cell composition and clonal organization. These findings extend a descriptive T-cell atlas by linking transcriptional phenotype to clonal expansion, anatomic distribution, and lesion-level composition within a paired study design.

The enrichment and clonal expansion of Tc17 cells reinforce a growing body of work positioning CD8+ type 17 states within psoriatic inflammation. Liao and colleagues previously identified two IL17A-expressing Tc17 subsets in psoriatic skin, including states with cytotoxic, type 1, and type 22 features, and used shared TCR sequences to propose relationships among activated, cytotoxic, and Tc17 populations ^5^. Other studies have documented overrepresentation of IL-17A- and IL-22-producing CD8+ T cells in lesions, persistence of psoriasis-associated IL-17-producing alpha-beta T-cell clones after clinical resolution, and a cutaneous transcriptional gradient that constrains Th17 and Tc17 identity ^6,7,11^. Our data extend these observations by quantifying Tc17-associated expansion across matched lesional, contralesional, non-lesional, and blood compartments. However, neither shared clonotypes nor RNA-velocity vectors establish the direction of differentiation. The observed relationships are best interpreted as compatible transcriptional and clonal connections among activated, memory, proliferating, and Tc17-associated states.

The paired sampling design also clarifies the anatomic organization of the TCR repertoire. Clonotype overlap was substantially greater between lesional and contralesional sites than between skin and blood, consistent with a repertoire that is strongly shaped by the cutaneous inflammatory environment. At the same time, the detection of shared blood–skin clonotypes indicates that tissue restriction is not absolute. Notably, cutaneous IL-17-associated clonotypes mapped to several circulating CD4+ and CD8+ phenotypes rather than to a single blood population. This result cautions against assigning one circulating state as the direct precursor of a cutaneous effector population from cross-sectional data.

Instead, it supports the broader view, emphasized in prior single-cell work from the Liao laboratory, that joint analysis of circulating and tissue compartments can reveal clonotype relationships that are obscured when either compartment is studied alone ^10^. The limited overlap with blood also helps explain why blood-only immune profiling may incompletely represent the lesion-level T-cell response.

Skin T cells expressed tissue-residency-associated programs across multiple phenotypic states. These programs were enriched for canonical tissue-retention genes and depleted for recirculation-associated genes, consistent with established human tissue-resident memory signatures ^14^. In psoriasis, CD103+ IL-17-competent resident-memory populations have been detected in active, resolved, and clinically uninvolved skin, and persistent local clones have been proposed to contribute to disease recurrence ^7,15,22^. Our observations are consistent with tissue adaptation across several skin T-cell states, but transcriptional signatures alone do not demonstrate long-term residence. Direct migration, retention, or longitudinal lineage-tracing experiments would be required to distinguish stable residents from recently recruited cells that have acquired a tissue-associated program.

The Treg findings should similarly be interpreted as state differences rather than as a direct measure of regulatory function. Lesional skin contained more Tregs and a shift toward states expressing TIGIT, CTLA4, ICOS, and other activation-associated markers. Earlier psoriasis studies described impaired Treg suppression and plasticity toward IL-17A-producing states ^12,13^. Joint work from the Rosenblum and Liao laboratories subsequently identified CD27 and OX40 as constraints on Th17-like differentiation in Tregs, while longitudinal profiling showed that cutaneous Treg transcriptional programs change during IL-17A blockade ^8,9^. The activated marker profile in our dataset is therefore biologically plausible, but it does not establish whether lesional Tregs are more suppressive, less suppressive, or functionally redirected. Functional assays and lineage-resolved analyses will be needed to distinguish altered abundance from altered regulatory activity.

Beyond individual cell states, the data identify recurrent patterns of lesion-level T-cell composition. Prior bulk-transcriptomic and spatial studies have demonstrated substantial molecular heterogeneity among psoriasis lesions and have described distinct inflammatory networks or multicellular ecosystems ^16-18^. Our two groups are narrower in scope: they were derived only from T-cell composition and should be considered compositional states rather than clinical or etiologic psoriasis subtypes. Their high concordance across paired lesions suggests that they capture a reproducible within-participant feature in this cohort. Nevertheless, the number of participants is modest, the state boundaries were derived within the discovery dataset, and independent validation will be required before these states can be linked to prognosis or therapeutic selection.

HLA-C*06:02 provides a biologically relevant context for the observed CD8+ architecture. The allele alters the class I peptide repertoire available for T-cell recognition and is the strongest common genetic risk factor for psoriasis ^2,20^. Structural and functional studies have begun to define how psoriasis-associated peptides can engage HLA-C*06:02-restricted TCRs ^4,21^. In our cohort, carrier status was associated with differences in selected CD8+ populations and in patterns of clonotype expansion and overlap. These observations are consistent with genotype-associated organization of the CD8+ response, but they do not identify the relevant antigens, demonstrate HLA restriction of the expanded clonotypes, or establish a causal effect of genotype. Paired TCR–peptide–HLA validation will be necessary to connect these descriptive associations to antigen specificity.

The longitudinal observations after therapy are deliberately exploratory. Prior work from Liao, Rosenblum, and colleagues showed that IL-17A blockade resolves most psoriasis-associated transcriptional differences in whole lesions and sorted cutaneous T-cell subsets while leaving selected residual programs ^9^. More recent single-cell studies have mapped the effects of IL-23 blockade on inflammatory T-cell, stromal, myeloid, and vascular circuits at substantially greater longitudinal depth ^23-25^. Our treated cohort included only eight participants receiving heterogeneous therapies. The variable changes in IL-17-associated cells, Tregs, clonotypes, and compositional-state assignments should therefore be viewed as within-cohort observations that motivate future longitudinal studies, not as estimates of treatment-specific mechanisms or predictors of response.

Several limitations define the appropriate scope of these conclusions. The study is observational, and cell recovery from dissociated biopsies may alter measured proportions. Multiple skin samples from the same participant improve within-participant comparisons but require careful attention to the participant as the unit of inference. RNA velocity, transcriptional tissue-residency scores, and clonotype sharing provide complementary evidence about relationships among states, but none directly establishes lineage direction, migratory behavior, antigen specificity, or cellular function. The treatment and HLA-C*06:02 analyses are limited by subgroup size, and the lesional compositional states were not tested in an independent cohort. Finally, transcript abundance does not establish cytokine secretion, and marker-defined Treg activation does not establish suppressive activity. These limitations motivate orthogonal validation using spatial methods, functional assays, peptide–HLA testing, and independent longitudinal cohorts.

In summary, paired analysis of multiple skin sites and blood links the transcriptional diversity of psoriasis T cells to their clonal and anatomic organization. The data support a model in which lesions contain expanded, tissue-adapted IL-17-associated populations embedded within broader and recurrent T-cell compositional states, with measurable but limited connections to the circulation and with variation associated with HLA-C*06:02 status. This framework complements prior UCSF studies of Tc17 diversity, skin Treg specialization, and treatment-associated remodeling by showing how phenotype and clonotype are coordinated across tissues within the same participants.

## Methods

### Participant recruitment and ethics

Study participants were recruited from the dermatology clinics at the University of California, San Francisco. Participants with psoriasis provided written informed consent under UCSF Institutional Review Board protocol 18-26971. Adults were eligible for the cross-sectional psoriasis cohort if they were at least 18 years of age, had plaque psoriasis for at least 6 months with affected body surface area >5%, had at least one plaque on the trunk or extremities with area >10 cm², and had not received biologic therapy within 12 weeks, oral systemic therapy within 4 weeks, or topical therapy within 2 weeks before sampling (Supplementary Table 1). A longitudinal cohort initiating or changing treatment met the same clinical criteria but was not required to meet the pretreatment washout criteria. The study included 23 participants with psoriasis and 11 healthy donors; four healthy donors contributed skin and seven contributed blood. Nineteen participants with psoriasis contributed both skin and blood samples, and eight contributed matched pre- and post-treatment samples. Study and sample metadata are summarized in Table 1.

### Biospecimen collection and processing

For participants with psoriasis, peripheral venous blood and 5-mm punch biopsies were collected where available. The skin-sampling design comprised one lesional site, a second lesional site on the contralateral side of the body, and one non-lesional site. For the longitudinal cohort, skin and blood were collected at baseline and again after 8–14 weeks of treatment. Three participants received risankizumab, three received guselkumab, and two received phototherapy. Healthy skin was obtained from tissue discarded during surgery unrelated to inflammatory skin disease from adults with no history of autoimmune disease; healthy PBMC samples were obtained from seven blood donors.

Biopsies were transported in HypoThermosol medium (BioLife Solutions) and cryopreserved in CryoStor CS10 medium (BioLife Solutions) at −80 °C for one day before transfer to liquid nitrogen. Frozen skin was thawed in phosphate-buffered saline, trimmed to remove hair and subcutaneous adipose tissue, finely minced, and incubated overnight in 5% CO₂ in RPMI 1640 supplemented with Collagenase Type IV (LS004188; Worthington Biochemical), DNase I (DN25-1G; Sigma-Aldrich), 10% fetal bovine serum, 1% HEPES, and 1% penicillin–streptomycin. Single-cell suspensions were passed through a nylon filter, centrifuged, and counted. PBMCs were isolated from peripheral blood by Ficoll density-gradient centrifugation. Genomic DNA was isolated from PBMCs, and HLA class I typing was performed to four-digit resolution by the UCSF Immunogenetics Laboratory.

### Antibodies and flow cytometry

Biopsy-derived and PBMC suspensions were stained with Sytox Blue for viability and with anti-human CD45 and CD3 antibodies (BD Biosciences). For samples with single-cell protein measurements, 0.5 µg of each TotalSeq antibody (BioLegend) was included in the presort stain. The antibody-derived tag panel comprised CD279, CD134, CD69, CD223, CD152, CD44, CD25, CD62L, CD4, and CD8α. Viable CD45+CD3+ cells were sorted into MACS buffer and resuspended for loading on the 10x Genomics platform, typically in 30 µL at a maximum concentration of 1,000 cells/µL.

### Single-cell library preparation and sequencing

Sorted T cells were loaded on a 10x Genomics Chromium Controller using the 5′ V(D)J workflow with a target recovery of 10,000 cells. Reverse transcription, emulsion breakage, and cDNA cleanup were performed according to the manufacturer’s protocol. cDNA was preamplified for 12 cycles, and 5% of the preamplified material was reserved for TCR enrichment by two rounds of nested multiplex PCR using human TRAC- and TRBC-specific primers. Final gene-expression and TCR libraries were diluted to 20 nM and pooled at equimolar ratios. Libraries were sequenced on an Illumina NovaSeq with a target depth of 500 million reads per sample using 26-bp read 1, 98-bp read 2, and an 8-bp i7 index read.

### Sequence processing and quality control

Gene-expression libraries were aligned to the human GRCh38 reference genome with Cell Ranger v6.0.0 (10x Genomics). Antibody-capture libraries, where available, were processed with the corresponding feature-barcode reference. TCR libraries were processed with the Cell Ranger V(D)J pipeline and the human TCR reference. Transcriptomic quality control and downstream analysis were performed in Seurat v4.3.0. Genes detected in fewer than three cells were excluded.

Cells were required to map to >200 genes, contain <10% mitochondrial transcripts, and contain >5% ribosomal transcripts. Cells with dissociation-stress scores >2 standard deviations above the mean and UMI-count outliers defined by the 5% analysis criterion were excluded, as were non-T-cell contaminants. Downstream analyses retained cells with one productive TCRα chain and one productive TCRβ chain. After filtering, 205,635 skin-derived T cells spanning 24,782 detected genes and 350,310 blood-derived T cells were retained.

### Transcriptomic integration, clustering, and annotation

Skin and blood datasets were normalized in Seurat, and sequencing batches were aligned with Harmony. Nearest-neighbor graphs and graph-based clusters were generated in the aligned space using FindNeighbors and FindClusters, and uniform manifold approximation and projection (UMAP) was used for visualization. This analysis resolved 19 major skin T-cell clusters and 18 major blood T-cell clusters. Integrated skin–blood analysis resolved 22 phenotypic populations. Cluster identities were assigned using canonical transcript markers and antibody-derived tags where available.

To mitigate dropout of lineage-marker transcripts, cells with unambiguous CD4 or CD8 expression were assigned directly; otherwise, lineage was assigned when a cell shared a paired TCR clonotype with a cell having an unambiguous CD4 or CD8 assignment. Skin and blood CD4+ and CD8+ compartments were subsequently renormalized and reclustered to resolve detailed cell states. Conventional CD4+ T cells and regulatory T cells were analyzed separately. For the integrated skin–blood analysis, differential-expression counts were summarized within each annotated population by the direction of the skin-versus-blood contrast.

### Tissue-residency-associated transcriptional analysis

Tissue-resident memory T-cell (TRM) programs were evaluated using a gene-signature score based on canonical tissue-retention and recirculation-associated markers. These included ITGAE and CXCR6 as tissue-retention markers and S1PR1 and SELL as recirculation markers. Signature scores and the corresponding TRM-like and non-TRM-like classifications were summarized across integrated cell states and tissue groups.

### TCR clonotype and sharing analyses

Productive paired TCR calls were integrated with the single-cell expression data using scRepertoire v1.7.2. Clonotypes were defined by exact matching of paired TCRα- and TCRβ-chain sequences. Cells were assigned to clonal-size categories according to the number of cells bearing the same clonotype: single (1 cell), small (2–5 cells), medium (6–20 cells), large (21–100 cells), and hyperexpanded (>100 cells). Clonal-size distributions were summarized by tissue and annotated cell state.

Clonotype overlap among tissue compartments and transcriptional states was quantified using the Morisita overlap index implemented in scRepertoire. Cluster-level heat maps displayed Morisita values >0.02 for clarity. Shared clonotypes were also used to map IL-17-associated skin cells to annotated blood populations. These analyses measured repertoire overlap and were not used to infer migration direction or precursor–product relationships.

### Cell-composition and embedding-density analyses

The abundance of each annotated T-cell state was calculated within each sample before group-level comparison. Differential abundance between lesional and non-lesional psoriasis skin, and between HLA-C*06:02 groups where indicated, was evaluated with the DCATS R package. Reported false-discovery-rate (FDR) values were used to account for multiple comparisons, with FDR <0.05 considered significant. To visualize shifts in cell-state distributions across the UMAP, Gaussian kernel densities were estimated separately by sample group with scanpy.tl.embedding_density; density values were scaled from 0 to 1 within each group.

### Lesional compositional-state analysis

For this analysis, both primary lesional and contralesional biopsies were considered lesional samples. A separate de novo 24-cluster solution was used for the compositional analysis. The abundance of each cluster was percentile-ranked across 57 lesional samples from 23 participants. Principal-component analysis was used to identify 14 clusters that contributed most strongly to separation among samples. The resulting 14-cluster composition profiles defined two groups, labeled compositional state 0 and compositional state 1. State assignments were compared descriptively across paired body sites and, where available, before and after treatment.

### HLA-C*06:02 analysis

Participants with at least one HLA-C*06:02 allele were classified as carriers (n = 8), and those without the allele were classified as non-carriers (n = 15). The genotype analysis was restricted to pre-treatment CD8+ T cells from lesional skin. CD8+ T-cell composition was compared between groups using the sample-level differential-abundance framework described above. Clonal-size distributions and Morisita overlap among selected CD8+ states were then summarized by HLA-C*06:02 status. TCR variable-gene usage and the compositions of conventional CD4+ and regulatory T-cell states were also summarized descriptively by carrier status.

### Longitudinal treatment analysis

Eight participants contributed matched baseline and post-treatment samples. For each participant, annotated cell-state frequencies, the ratio of IL-17-associated T cells to Tregs, and clonotype detection across time points were summarized. Clonotypes were categorized as detected only before treatment, at both time points, or only after treatment. Because the cohort was small and included three treatment modalities, longitudinal results were treated as descriptive and were not used for treatment-specific mechanistic inference.

### RNA-velocity analysis

Cell Ranger-aligned BAM files were converted to loom files containing spliced and unspliced transcript counts using velocyto CLI v0.17.17. Loom files were merged with the annotated T-cell objects, normalized, and filtered. RNA velocity was estimated with the dynamical model in scVelo, and velocity stream plots were projected onto the UMAP embedding. MAIT cells were excluded from the CD8+ velocity visualization.

## Figure Legends

**Figure S1.**
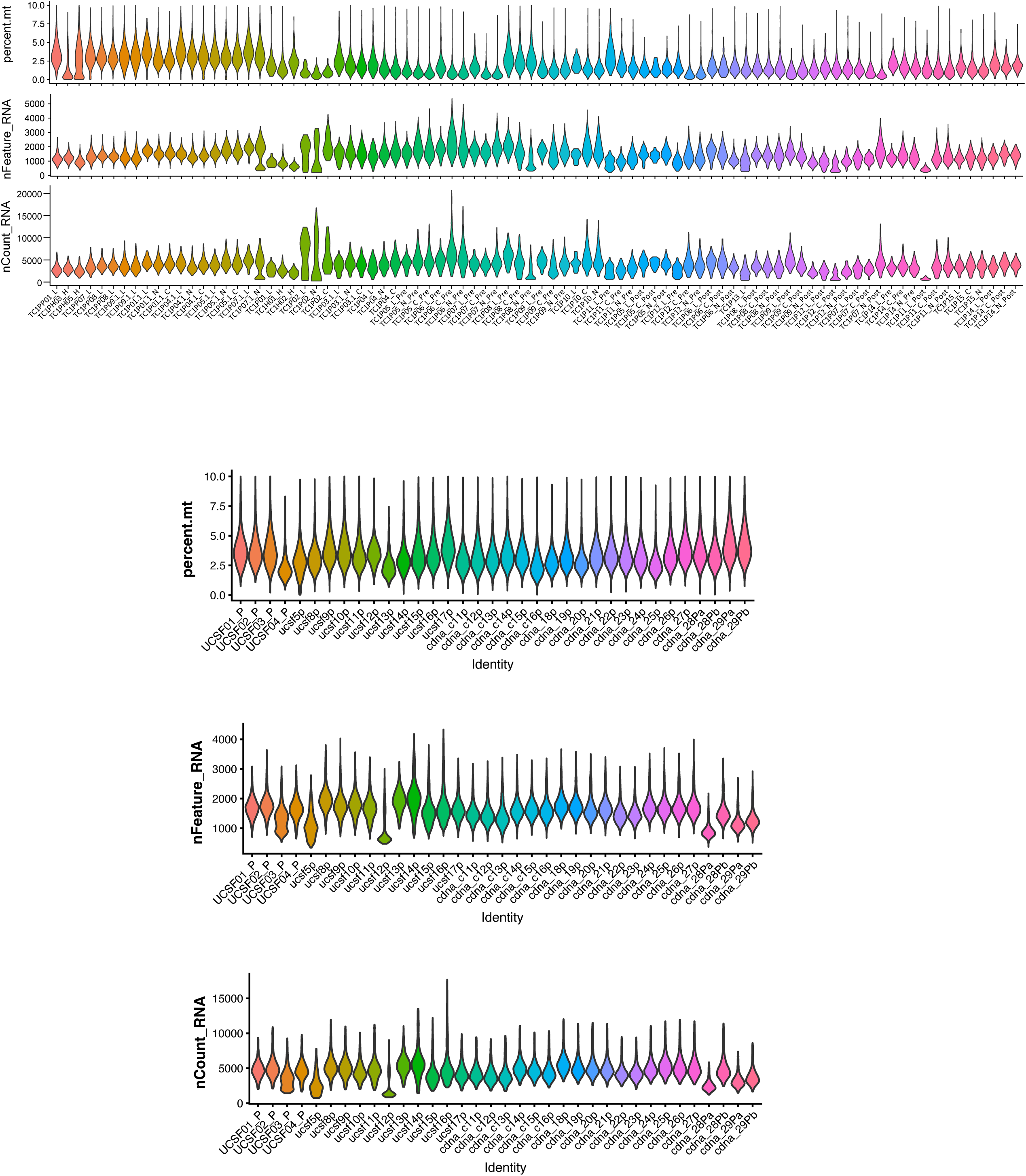
Single-cell RNA-sequencing quality-control distributions. Violin plots show the percentage of mitochondrial transcripts (percent.mt), number of detected genes (nFeature_RNA), and total UMI count (nCount_RNA) for each sequencing library. Skin-derived libraries are shown in the upper set of plots and PBMC-derived libraries in the lower set. Each violin summarizes the per-cell distribution within one library.

**Figure S2.**
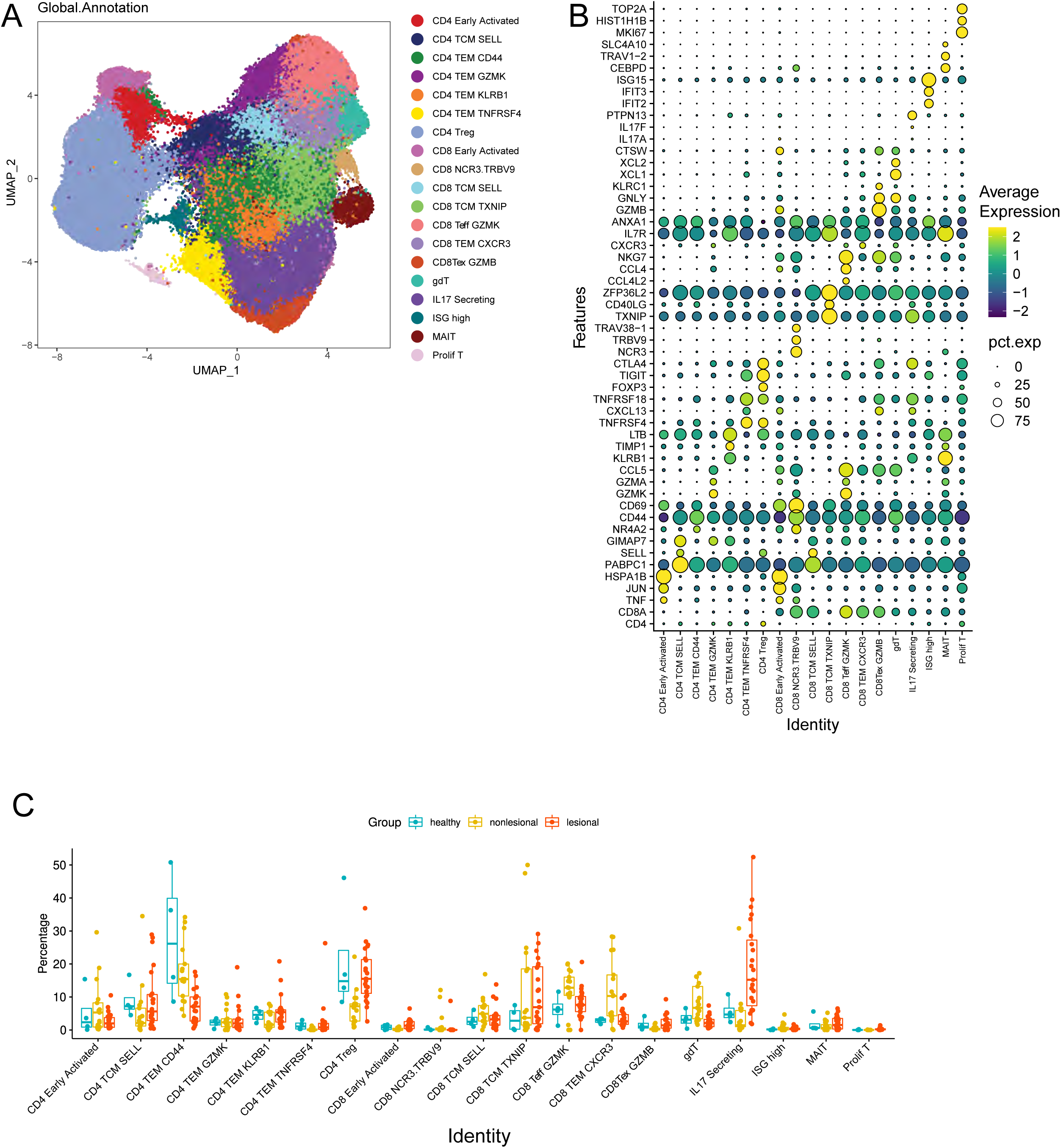
Global skin T-cell landscape including healthy skin. (A) UMAP of skin-derived T cells colored by global annotation. (B) Dot plot of marker-gene expression across the global states; dot size indicates the percentage of cells expressing each gene, and color indicates scaled average expression. (C) Sample-level frequencies of the annotated states in healthy skin (n = 4 samples), pretreatment non-lesional psoriasis skin (n = 17 samples), and pretreatment lesional psoriasis skin (n = 23 samples). Each point represents one sample; boxes summarize the median and interquartile range.

**Figure S3.**
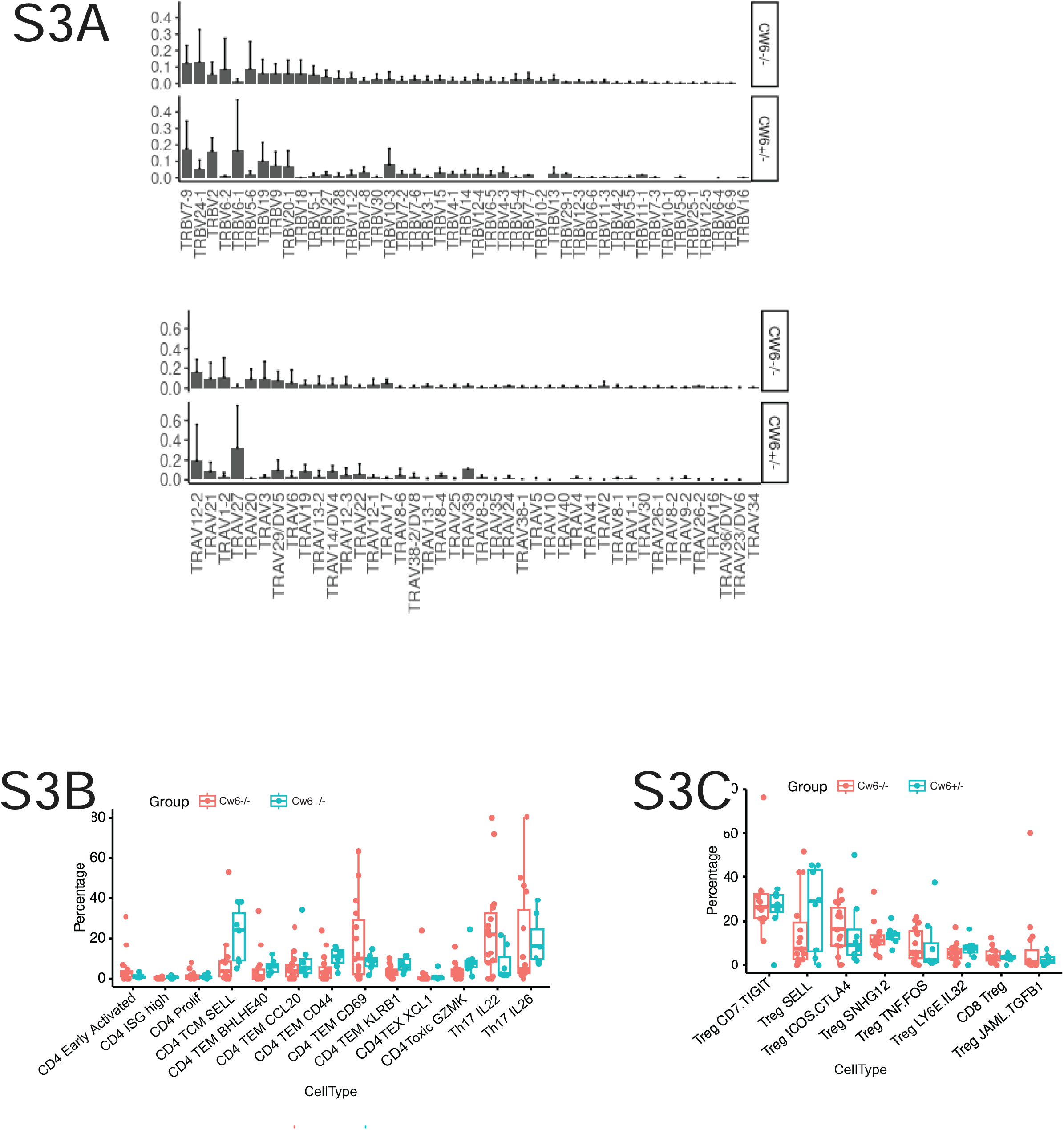
HLA-C*06:02-stratified supplemental analyses. (A) TRBV and TRAV usage in lesional CD8+ T cells. (B) Conventional CD4+ T-cell frequencies. (C) Treg frequencies. Points are participants; boxes show the median and interquartile range.

## Acknowledgements

We would like to thank Edward Hadeler, Mimi Chung, Erin Bartholomew, Julie Hong, Megan Mosca, Nicholas Brownstone, and Sugandh Kumar for their technical assistance with this project.

## Supplementary Tables

**Supplementary Table 1.**
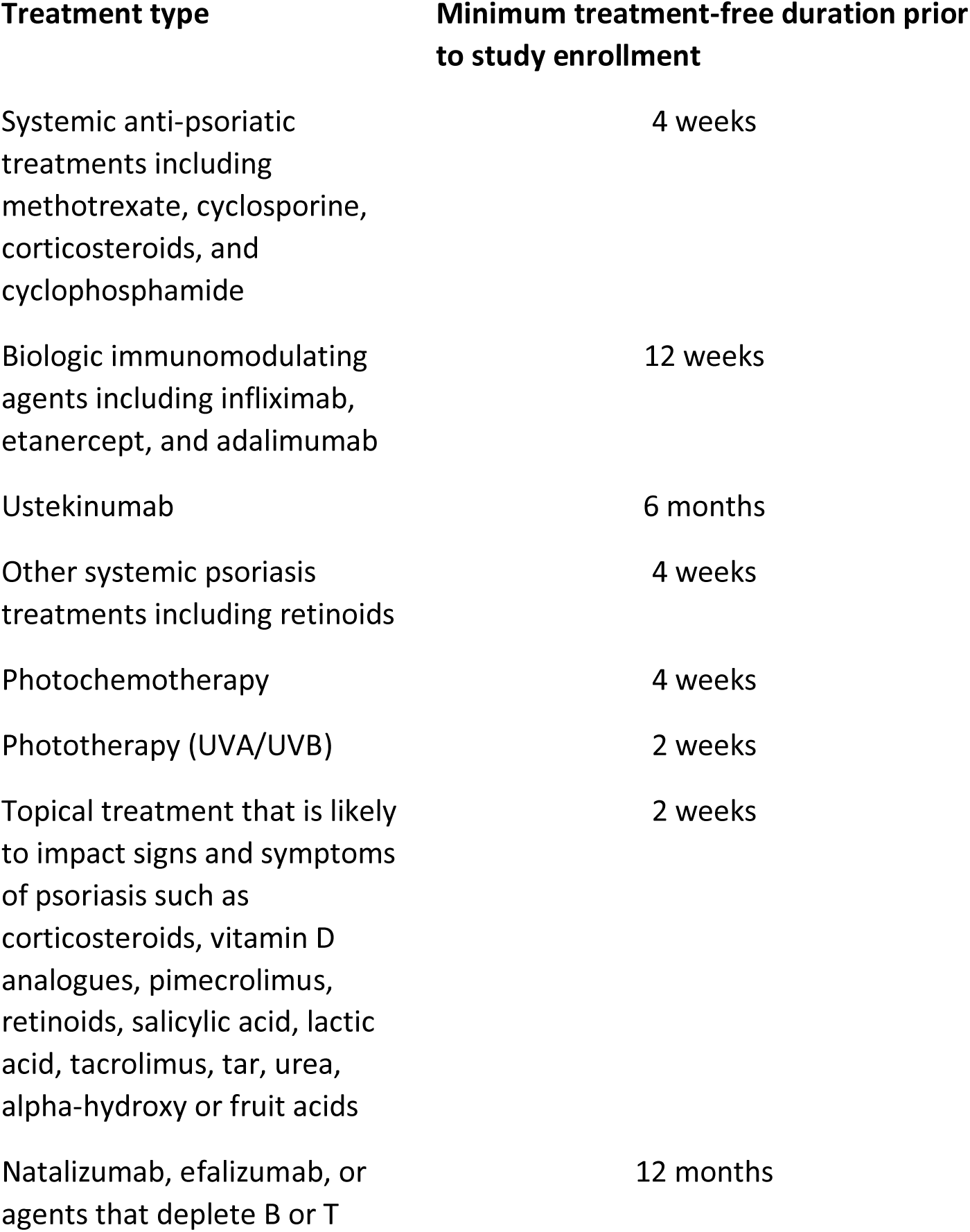
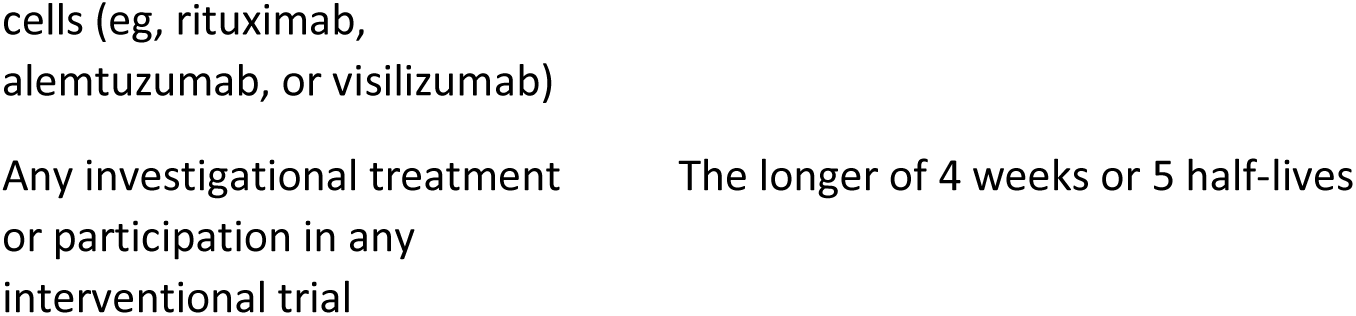
Prohibited Medications and Treatments for Patient Enrollment.

